# Degradation of the incretin hormone Glucagon-Like Peptide-1 (GLP-1) by *Enterococcus faecalis* metalloprotease GelE

**DOI:** 10.1101/732495

**Authors:** Stephanie L. LeValley, Catherine Tomaro-Duchesneau, Robert A. Britton

## Abstract

Metabolic diseases, including Type 2 Diabetes and obesity, have become increasingly prevalent global health concerns. Studies over the past decade have established connections between the gastrointestinal microbiota and host metabolism, but the mechanisms behind these connections are only beginning to be understood. We were interested in identifying microbes that have the ability to modulate the levels of the incretin hormone glucagon like peptide 1 (GLP-1). Using a human derived cell line that is capable of secreting GLP-1 in response to stimulatory ligands (NCI-H716), we identified supernatants from several bacterial isolates that were capable of decreasing GLP-1 levels, including several strains of *Enterococcus faecalis*. We further identified the secreted protease GelE, an established virulence factor from *E. faecalis*, as being responsible for GLP-1 inhibition via direct cleavage of GLP-1 by GelE. Finally, we demonstrated that *E. faecalis* supernatants can disrupt a colonic epithelial monolayer and cleave GLP-1 in a *gelE* dependent manner. This work suggests that a secreted factor from an intestinal microbe can traverse the epithelial barrier and impact levels of an important intestinal hormone.

**Importance:** Humans have a complex and interconnected relationship with their gastrointestinal microbiomes, yet our interest in the microbiome tends to focus on overt pathogenic or probiotic activities, leaving the roles that commensal species may have on host physiology and metabolic processes largely unexplored. Commensal organisms in the microbiome produce and secrete many factors that have an opportunity to interact with the gastrointestinal tract and host biology. Here we show that a secreted protease from *E. faecalis*, GelE, is able to degrade the gastrointestinal hormone GLP-1, which is responsible for regulating glucose homeostasis and appetite in the body. The disruption of natural GLP-1 signaling by GelE may have significant consequences for maintaining healthy blood glucose levels and in the development of metabolic disease. Furthermore, this work deepens our understanding of specific host-microbiome interactions.

## Introduction

The human gastrointestinal (GI) tract is home to trillions of microorganisms, collectively referred to as the GI microbiome (1). Because of the direct physical proximity that the gut microbiome has with its human host, it is no surprise that the microbiome plays a role in multiple aspects of health, the best characterized of these being immune tolerance, pathogen resistance, and digestion (2). Less understood are the interactions between the microbiome and human metabolism. Despite limited mechanistic insight into the cross-section of microbiome and host metabolism, it is of great interest as both an etiology of disease and for potential therapeutic applications, and some understanding is beginning to emerge. The first insights into microbial influence over human metabolism came from studies demonstrating that the simple absence of a microbiome resulted in decreased total body fat in germ-free mice, compared to conventional mice, independent of food intake; further, the decrease in fat mass could be gained back by colonizing the germ-free mice with bacterial communities from conventionally raised mice (3). The observed ability of the microbiome to help harvest energy from the diet sparked a variety of research studies over the next decade, with a focus on the interaction of the microbiome with GI hormone peptides, in particular the nutrient-stimulated incretin Glucagon-like Peptide-1 (GLP-1). Already a therapeutic target for Type 2 Diabetes (T2D), GLP-1 is an integral signaling hormone responsible for promoting insulin secretion and satiety, while decreasing glucagon secretion and gastric emptying. Early work showed that addition of the prebiotic oligofructose to the diets of rats on a high-fat diet increased GLP-1 levels measured from the portal vein, in addition to protecting from weight gain (4). Additional studies demonstrated that administration of *Akkermansia muciniphila* could reverse high-fat diet induced metabolic disorders, and that this activity was mediated at least partially by an outer membrane protein purified from *A. muciniphila* interacting with Toll-like receptor 2 (5, 6). These studies strikingly demonstrate that a specific bacterial species and factor are capable of impacting metabolic disease phenotypes. Despite a heightened research interest, additional mechanisms behind how the microbiome and host metabolism influence each other still remain largely undescribed.

Some of the obvious suspects to investigate for host-microbiome interactions are the many secreted proteins and metabolites that bacteria release as part of their natural lifecycle. While these external products are often part of bacterial cellular metabolism or provide beneficial function to the bacterial cell, they also have an opportunity to interact with their environment, in this case the human GI tract. Here, our screening for bacterial modulators of GLP-1 revealed multiple bacterial strains that can inhibit GLP-1 levels in an in vitro assay. We further characterize this inhibition as direct cleavage from *E. faecalis* strains by its secreted protease GelE, revealing a novel cleavage target of GelE. Finally, we suggest a role for GelE in disrupting natural GLP-1 signaling and metabolic processes.

## Results

### Screening a human-derived bacterial library for GLP-1 modulatory activity

Bacterial strains capable of modulating GLP-1 levels were identified by an in vitro screening pipeline using the GLP-1 secreting human cell line NCI H716 (7). Over 1500 cell-free supernatants collected from individual bacterial isolates were prepared and applied to NCI H716 cell monolayers for 2 h, and secretion of GLP-1 into the medium was measured by ELISA. NCI H716 cell viability was also monitored by PrestoBlue Cell Viability Reagent to ensure no significant increase in NCI H716 cell lysis or death (data not shown). The majority of bacterial isolates screened had no impact on GLP-1 levels; however, approximately 20 isolates showed a marked decrease in GLP-1 levels, many of them below the limit of detection of the ELISA (**Figure 1A**). We also identified 45 isolates that dramatically increase GLP-1 levels; these were further characterized in a separate study (8).

**Figure 1.**
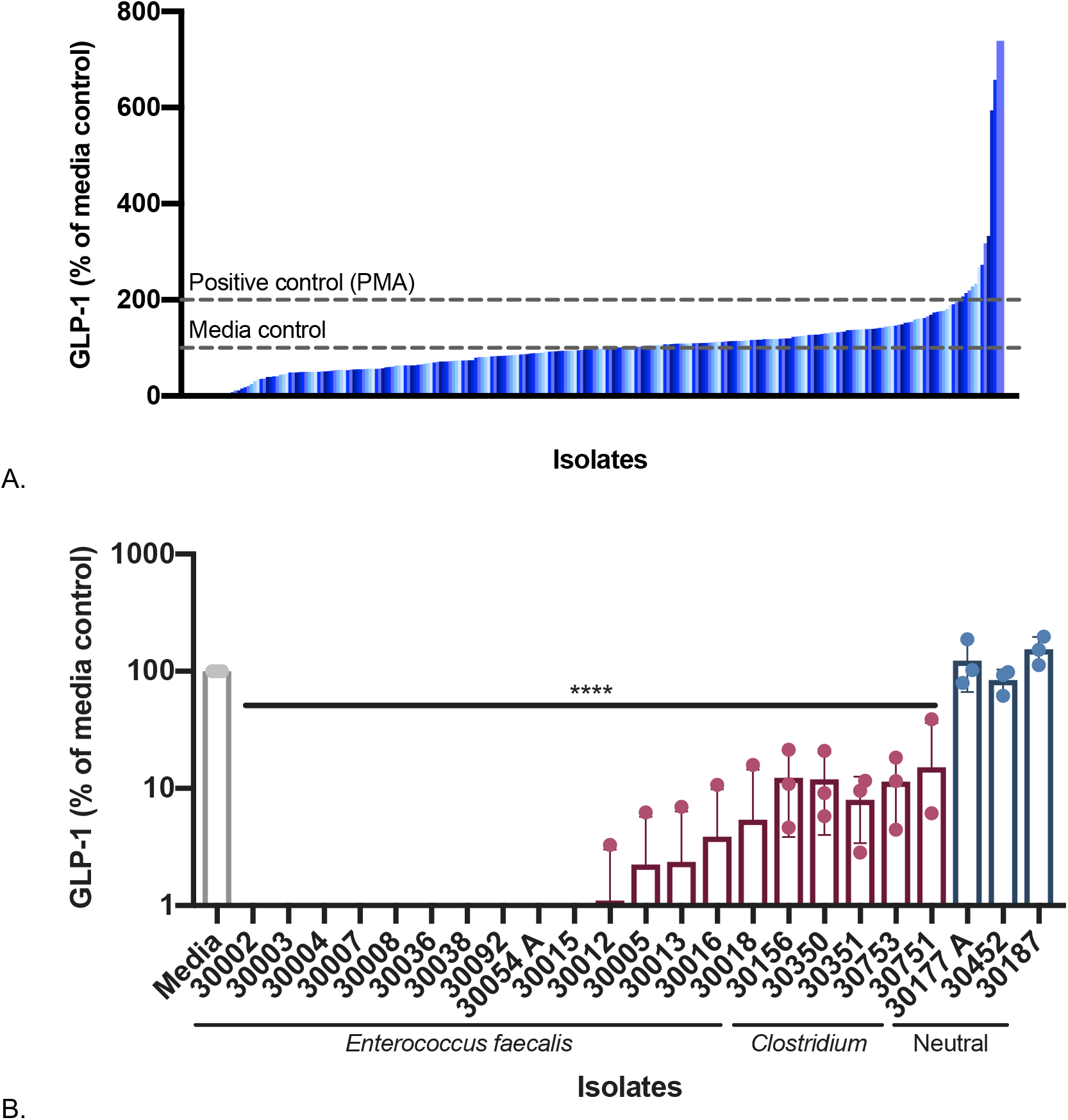
Screening for modulation of GLP-1. Bacterial supernatants have a wide range of effects on GLP-1 secretion from NCI H716 cells (A). Multiple isolates of *Enterococcus faecalis* and *Clostridium* decreased GLP-1 levels compared to media control (B). Data were obtained from three independent experiments (n = 3) and expressed as mean values ± SD (****p < 0.0001).

To identify the species of each isolate, the 16S rRNA gene was sequenced. The majority of the isolates identified as *Enterococcus faecalis*, as well as *Clostridium perfringens, C. bifermentans*, and *C. butyricum*. The *E. faecalis* isolates exhibited a stronger GLP-1 inhibitory effect, ranging between 0% ± 0.0 and 5.44 ± 9.1% GLP-1, compared to media controls (p < 0.0001, **Figure 1B**). The *Clostridium* species, while still inhibitory, consistently show a slightly weaker inhibitory effect, ranging between 8.0% ± 4.6 and 15.2% ± 21.0 GLP-1, compared to media controls (p < 0.0001, **Figure 1B**). Because of this difference in the GLP-1 inhibitory activities of *E. faecalis* and *Clostridium* species, we decided to further characterize the activity of the *E. faecalis* isolates.

### Identifying the factor secreted from E. faecalis responsible for GLP-1 inhibition

Size fractionation experiments showed that GLP-1 inhibitory activity from *E. faecalis* was contained within the 30-50 kDa size fraction of supernatant (**Figure S1**). We also found that the GLP-1 degradation activity was sensitive to heat and the metal ion chelator N,N,N’,N’-tetrakis(2-pyridylmethyl)ethane-1,2-diamine (TPEN) (**Figures S2 and S3**). These data suggest the factor responsible for GLP-1 inhibitory activity is a secreted, metal-dependent protein. Two secreted proteases from *E. faecalis* are well known, the serine protease SprE and the metalloprotease GelE, both of which are regulated by the two-component, quorum-sensing *fsr* (*faecalis* system regulator) operon (9) (**Figure 2A**). We obtained strains of *E. faecalis* containing null mutations in *sprE* (TX5243), *gelE* (TX5264), *fsrB* (TX5266), and a *gelE, sprE* double mutant (TX5128), all generated in the commonly used wild-type OG1RF *E. faecalis* strain (10). To test the GLP-1 inhibitory activity of these strains, supernatants were incubated directly with recombinant GLP-1 (Tocris) (**Figure 2B**). Of the strains tested, only the *DsprE* strain maintained GLP-1 inhibitory activity equal to that of wild-type OG1RF (OG1RF −0.06% ± 0.76, Δ*sprE* −0.14% ± 0.74, *p* < 0.0001), while the Δ*gelE*, Δ*fsrB*, and Δ*gelE*;Δ*sprE* strains no longer showed decreased GLP-1 levels (95.19% ± 4.48, 97.8% ± 7.37, 88.41% ± 17.47, respectively). Taken together, the loss of GelE, either directly by knock-out or indirectly by dysregulation through FsrB, results in a loss of GLP-1 inhibitory activity, suggesting direct cleavage of GLP-1 by the metalloprotease GelE.

**Figure 2.**
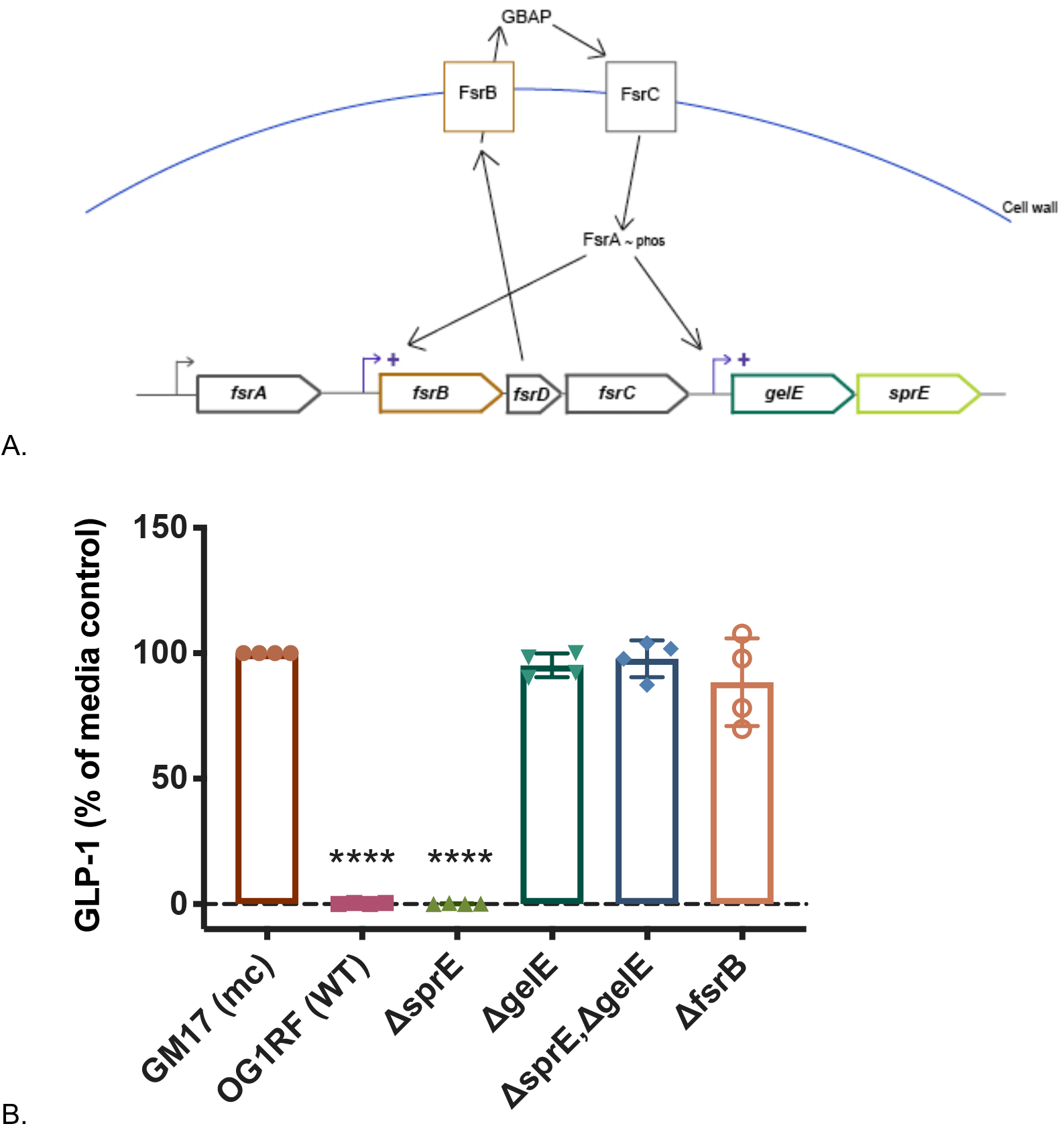
GelE is the factor responsible for GLP-1inhibition. Genetic pathway for production of *E. faecalis* proteases *gelE* and *sprE* (A). Activation of the quorum sensing system *fsr* activates transcription of *gelE* and *sprE*, immediately downstream of the *fsr* operon. *E. faecalis* strains unable to produce a functional GelE no longer deplete GLP-1 levels in an in vitro co-incubation of bacterial supernatants with GLP-1 (B). Data were obtained from four independent experiments (n = 4) and expressed as mean values ± SD (****p < 0.0001). GM17 (mc), bacterial culture media control.

To further characterize GLP-1 cleavage by GelE, the expression of *gelE* in five *E. faecalis* isolates was measured by quantitative PCR in relation to their ability to cleave GLP-1 (**Figure 3**). During the initial screen, we identified one *E. faecalis* isolate that did not inhibit GLP-1, a clinical isolate *E. faecalis* S613 (from Cesar A. Arias’s laboratory, University of Texas, Health Science Center) (11). The four GLP-1 degrading isolates highly express *gelE*, with expression levels at least 10-fold higher and up to 380-fold higher than the non-GLP-1 degrading isolate S613, demonstrating a correlation between *gelE* expression and GLP-1 degradation.

**Figure 3.**
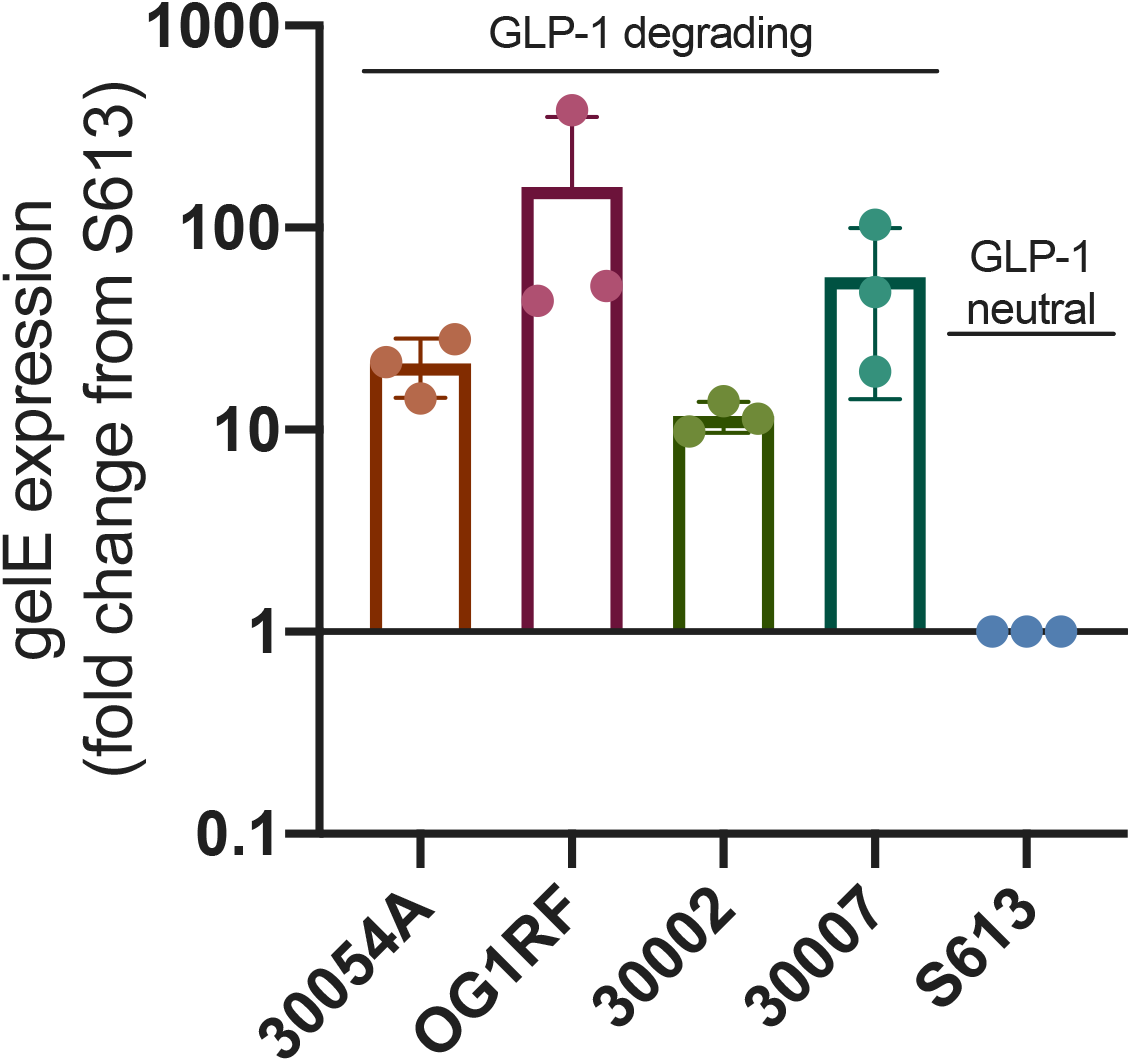
Expression of gelE correlates with GLP-1 degradation. GLP-1 degrading strains of *E. faecalis* have significantly higher expression levels of *gelE* compared to a GLP-1 neutral strain S613. Data were obtained from three independent experiments (n = 3).

### GelE specificity for human metabolic substrates

Previously, it has been shown that GelE degrades a range of substrates (12); thus, we wanted to better understand the range and specificity of GelE cleavage targets relevant to GLP-1 and other proteins involved in human metabolism. Supernatants from *E. faecalis* strains 30054A, 30002, OG1RF, Δ*gelE*, Δ*sprE*, and S613 were incubated with a panel of recombinant protein substrates (GLP-1, Glucose-dependent insulinotropic peptide (GIP), Peptide YY (PYY), leptin, glucagon, pancreatic peptide, insulin, IL-6, tumor necrosis factor alpha (TNFα), and monocyte chemoattractant protein-1 (MCP-1)) and quantified by a Luminex assay. From our findings, GelE is capable of degrading, to some extent, nearly all the substrates tested in the metabolic panel (**Figure 4**); however, some patterns of degradation emerged. Recapitulating our previous findings, GLP-1 levels were reduced to 0.57% ± 0.13, down to the limit of detection of the assay. Similar degradation was observed for GIP, glucagon, leptin, PYY, and pancreatic peptide. For insulin, MCP-1, and TNFα, degradation was less striking, but still statistically significant. Finally, we did not observe consistent degradation of IL-6 by GelE.

**Figure 4.**
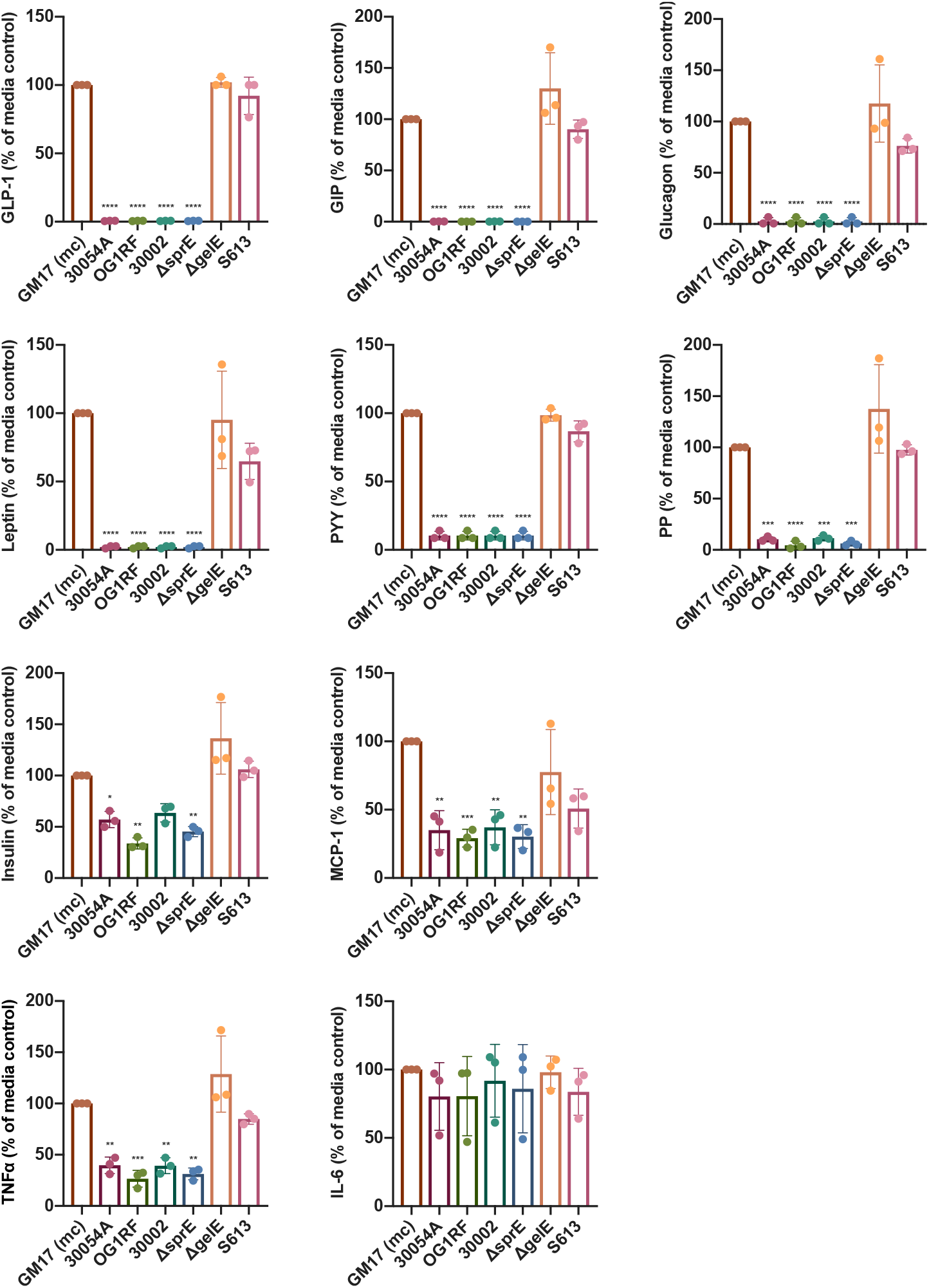
GelE specificity for human metabolic substrates. Data were obtained from three independent experiments (n = 3) and expressed as mean values ± SD (GLP-1, GIP, Glucagon, Leptin, PYY, and PP: ****p < 0.0001; Insulin: 30054A *p = 0.0351, OG1RF *p = 0.001, ΔsprE **p = 0.0058; MCP-1: 30054A **p = 0.0021, OG1RF ***p = 0.0009, 30002 **p = 0.0029, ΔsprE **p = 0.0011; TNFα: 30054A **p = 0.0038, OG1RF ***p = 0.0006, 30002 **p = 0.0036, ΔsprE **p = 0.0012). GM17 (mc), bacterial culture media control.

To gain a better understanding of the substrate preference of GelE, we also tested the ability of diluted supernatants of *E. faecalis* 30054A, from 1-0.0001X, to degrade the same panel of metabolic substrates (**Table 1**). Most prominently, supernatant from *E. faecalis* 30054A diluted to 0.01X still degraded the majority of GLP-1 present, leaving only 10.38% ± 3.52 GLP-1 remaining. GIP, glucagon, and leptin also show high degradation with diluted supernatants (1.55% ± 1.09, 3.08% ± 3.25, and 7.95% ± 4.72 substrate remaining, respectively, for 0.1X supernatants). This level of degradation does not hold true for all substrates tested: pancreatic peptide and PYY maintain fairly high levels of degradation with undiluted supernatants (10.35% ± 2.4 and 10.45% ± 3.1 substrate remaining, respectively, for undiluted supernatants), but degradation lessens upon dilution (47.49% ± 17.15 and 54.61% ± 13.14 substrate remaining, respectively, for 0.1X supernatants). Finally, MCP-1, TNFa, and insulin show less degradation even with undiluted supernatants (35.14% ± 14.1, 39.73% ± 8.0, and 57.29% ± 7.7 substrate remaining, respectively, for undiluted supernatants).

**Table 1:**
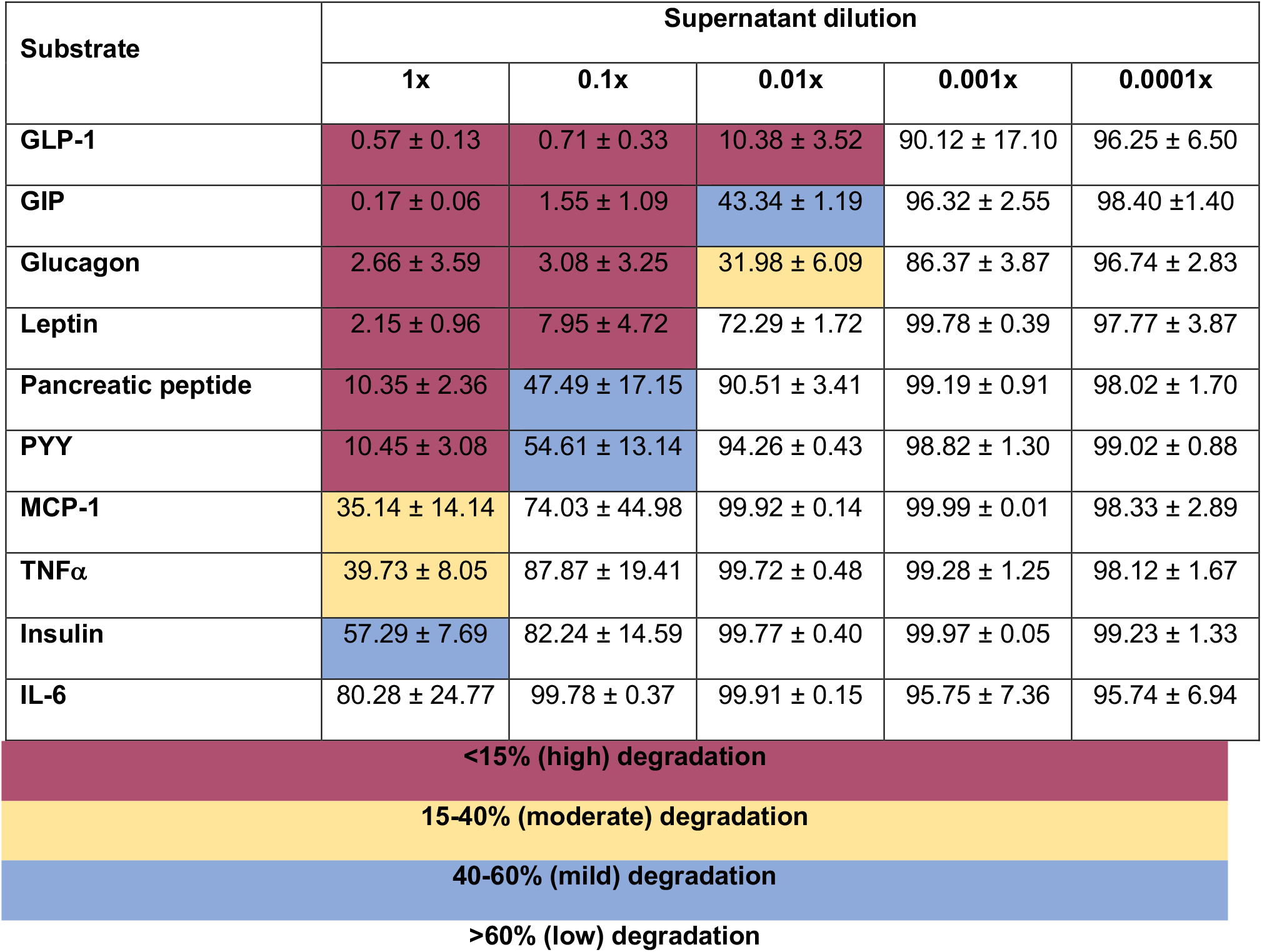
Degradation of metabolic substrates by E. faecalis 30054A diluted supernatant. Dilutions of supernatant from *E. faecalis* 30054A show an uneven distribution of degradation. Values represent the quantity of substrate remaining after incubation with supernatants. Data were obtained from three independent experiments (n = 3) and expressed as mean values ± SD.

| Substrate | Supernatant dilution |  |  |  |  |
| --- | --- | --- | --- | --- | --- |
|  | 1x | 0.1x | 0.01x | 0.001x | 0.0001x |
| GLP-1 | 0.57 ± 0.13 | 0.71 ± 0.33 | 10.38 ± 3.52 | 90.12 ± 17.10 | 96.25 ± 6.50 |
| GIP | 0.17 ± 0.06 | 1.55 ± 1.09 | 43.34 ± 1.19 | 96.32 ± 2.55 | 98.40 ± 1.40 |
| Glucagon | 2.66 ± 3.59 | 3.08 ± 3.25 | 31.98 ± 6.09 | 86.37 ± 3.87 | 96.74 ± 2.83 |
| Leptin | 2.15 ± 0.96 | 7.95 ± 4.72 | 72.29 ± 1.72 | 99.78 ± 0.39 | 97.77 ± 3.87 |
| Pancreatic peptide | 10.35 ± 2.36 | 47.49 ± 17.15 | 90.51 ± 3.41 | 99.19 ± 0.91 | 98.02 ± 1.70 |
| PYY | 10.45 ± 3.08 | 54.61 ± 13.14 | 94.26 ± 0.43 | 98.82 ± 1.30 | 99.02 ± 0.88 |
| MCP-1 | 35.14 ± 14.14 | 74.03 ± 44.98 | 99.92 ± 0.14 | 99.99 ± 0.01 | 98.33 ± 2.89 |
| TNFα | 39.73 ± 8.05 | 87.87 ± 19.41 | 99.72 ± 0.48 | 99.28 ± 1.25 | 98.12 ± 1.67 |
| Insulin | 57.29 ± 7.69 | 82.24 ± 14.59 | 99.77 ± 0.40 | 99.97 ± 0.05 | 99.23 ± 1.33 |
| IL-6 | 80.28 ± 24.77 | 99.78 ± 0.37 | 99.91 ± 0.15 | 95.75 ± 7.36 | 95.74 ± 6.94 |

|  |
| --- |
| <15% (high) degradation |
| 15-40% (moderate) degradation |
| 40-60% (mild) degradation |
| >60% (low) degradation |

### Interaction of GelE and GLP-1 through an epithelial layer

While GelE from *E. faecalis* may readily cleave GLP-1 and other substrates in vitro, it is important to consider the GI epithelium separating these two molecules in vivo. Previous work has implicated *E. faecalis*, and specifically GelE, in contributing to intestinal epithelium disruption (13–15). We aimed to model the ability of GelE to contact GLP-1 through an epithelial layer using T84 epithelial cells in a transwell format, to mimic the microbial interface on the apical side and the presence of GLP-1 on the basolateral side of the epithelium (**Figure 5A**). The integrity of the T84 epithelial layer, as measured by transepithelial electrical resistance, decreased by approximately half with the apical addition of cell-free *E. faecalis* supernatants expressing GelE, 30054A and OG1RF (52.2% ± 17.97, p = 0.0022 and 65.5% ± 9.47, p = 0.0439, respectively, compared to DMEM/F12 T84 cell culture media control); while *E. faecalis* supernatants lacking GelE, *DgelE* and S613, increased the integrity of the epithelial layer (156.6% ± 24.76, p = 0.0045 and 153.7% ± 18.93, p = 0.0084, respectively, compared to DMEM/F12 T84 cell culture media control) (**Figure 5B**).

**Figure 5.**
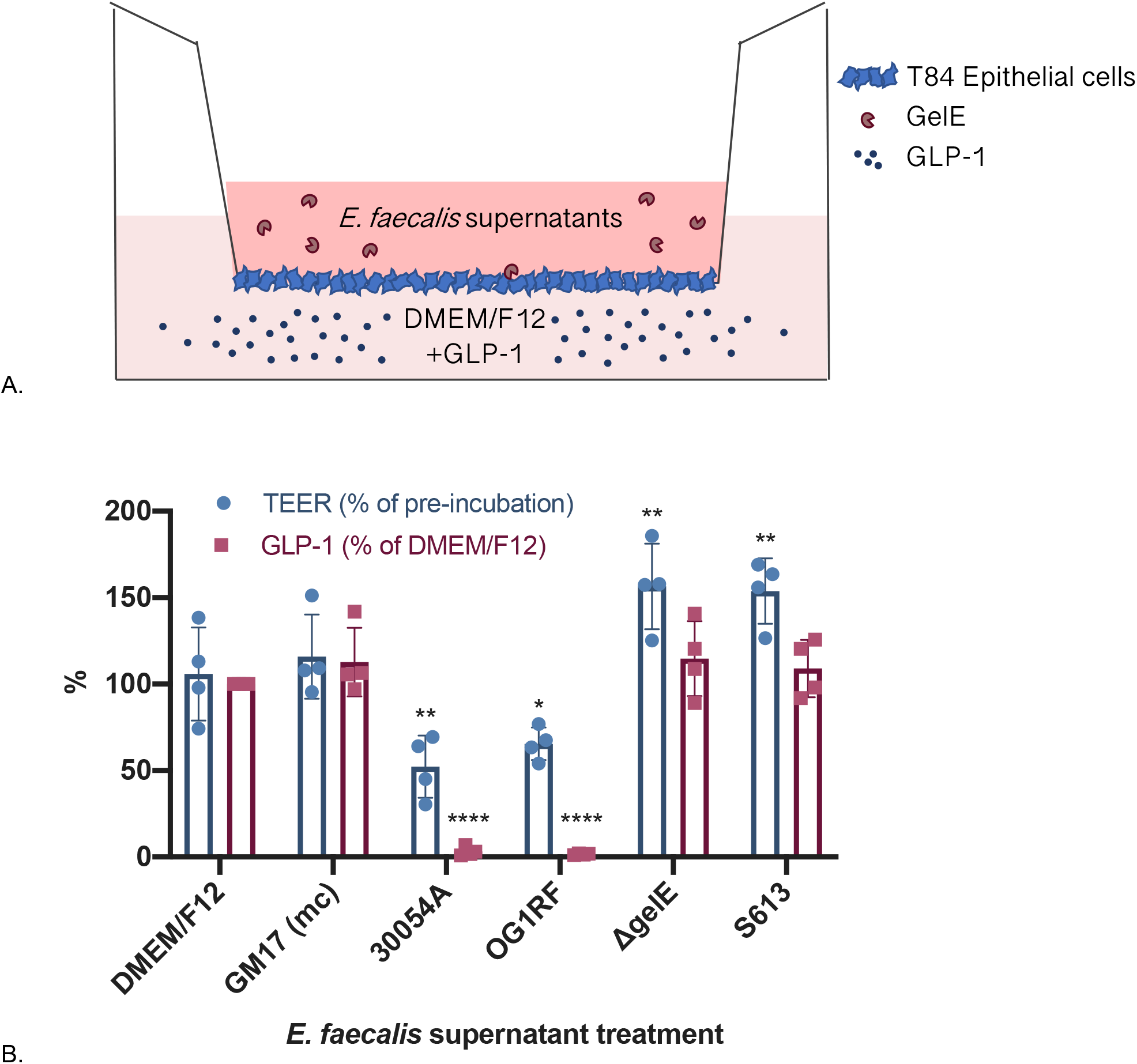
Degradation of GLP-1 through T84 epithelial layer. Schematic of experimental setup: T84 epithelial cells seeded onto transwell insert, with *E. faecalis* supernatants added to the upper chamber and GLP-1 in cell culture media (DMEM/F12) to the lower chamber (A) Schematic not drawn to scale. GelE producing supernatants decrease the integrity of a T84 epithelial cell layer while degrading GLP-1 on the basolateral side of the epithelium, compared to DMEM/F12 T84 cell culture media control (B). Data were obtained from four independent experiments and expressed as mean values ± SD (TEER: 30054A **p = 0.0022, OG1RF *p = 0.0439, ΔgelE **p = 0.0045, S613 **p = 0.0084; GLP-1: 30054A and OG1RF ****p < 0.0001). GM17 (mc), bacterial culture media control.

The basolateral compartment of each transwell contained approximately 500 pM GLP-1 supplemented into the DMEM/F12 media. When GelE containing supernatants 30054A or OG1RF were added apically to the T84 cell epithelial layer, GLP-1 in the basolateral compartment was cleaved nearly completely (3.1% ± 2.68, p < 0.0001 and 1.6% ± 0.46, p < 0.0001 respectively, compared to DMEM/F12 T84 cell culture media control), while GLP-1 in the basolateral compartment of apical supernatants from *DgelE* and S613 *E. faecalis* strains, remained intact (**Figure 5B**). Together, these data demonstrate that GelE causes moderate damage to an epithelial layer, allowing access to the basolateral side where it can cleave GLP-1.

## Discussion

Bacterial cells secrete a wide range of proteins and metabolites during their lifecycle, and one such class of secreted molecules are proteases, enzymes that break down other proteins or peptides. A well-studied example of this is *Enterococcus faecalis* and its gelatinase GelE. A secreted metalloprotease, GelE serves *E. faecalis* by degrading misfolded surface proteins and decreasing chain length for dissemination (16). A role for GelE in preventing biofilm formation has also been described by multiple research groups (17, 18).

Because *E. faecalis* has been implicated in various infections, including endocarditis, bacteremia, and urinary tract infections (19), GelE has been studied for its interaction with human proteins. A thorough characterization of GelE revealed multiple host cleavage targets, including glucagon and cholecystokinin, among several other substrates (12). More recently, GelE has been shown to cleave the C3-α chain of the human complement system, promoting immune evasion (20). Additionally, GelE can degrade the tight junction protein E-cadherin, contributing to intestinal inflammation and impaired barrier integrity (13). Our work adds GLP-1, among other metabolic factors, to the list of GelE targets.

Our data from Luminex assays using a panel of metabolic substrates show that GelE is able to degrade more substrates than previously suspected, and furthermore, that GelE has an enhanced ability to degrade some substrates over others, including GLP-1, GIP, glucagon, and leptin. If occurring in the human body, degradation of these various substrates could have differing, sometimes contradictory, effects on host metabolic processes. GLP-1, GIP, and PYY are secreted basolaterally by enteroendocrine cells and have beneficial functions for host metabolism, and their absence would likely permit a change in glucose homeostasis, leading to hyperglycemia, and an increase in food intake. Made and secreted in the pancreas, degradation of insulin by GelE would likely result in similar hyperglycemia, and a reduction in pancreatic peptide might allow for increased food intake. Conversely, degradation of glucagon from alpha cells of the pancreas might result in hypoglycemia. Leptin is produced in adipocytes, and its degradation would likely cause dysregulation of fat accumulation and processing. Finally, MCP-1 and TNFa are pro-inflammatory cytokines, and disruption of their signaling by degradation from GelE might delay an immune response during pathogen invasion. Importantly, whether these cleavage events are occurring and have relevant consequences in a whole biological system needs to be confirmed in animal studies. Based on proximity, GelE crossing the epithelial layer would first encounter molecules secreted into the lamina propria, and so we suspect GI hormones such as GLP-1, GIP, and PYY would be primary targets for degradation by GelE.

Our study supports the findings that *E. faecalis* and its protease GelE can compromise an epithelial layer and gain access to the basolateral environment (13–15). This is not surprising from a clinical perspective, as *E. faecalis* is implicated in cases of bacteremia and sepsis (23). For the topic of this study, this access could allow GelE to contact vital substrates responsible for host metabolic homeostasis as they are secreted into the GI lamina propria. Even mild inflammation often observed in individuals with metabolic syndrome, coupled with bacterial instigators like GelE from *E. faecalis*, could create a weakened epithelium (24). Once the integrity of the epithelium is damaged, luminal contents have an opportunity to move from the lumen of the GI tract and into the lamina propria just on the other side of the epithelial layer, where many hormones and metabolites are secreted before moving into the circulatory system. Furthermore, there is little indication regarding whether GelE could also be capable of diffusing into circulation as most hormones and nutrients do. Additionally, others have demonstrated by proof of concept that the microbiome encodes DPP-4 like activity that can traverse the epithelium, and further propose that this activity is capable of modulating protein digestion and ultimately host metabolism and behavior (25); while DPP-4 and GelE work via different proteolytic mechanisms, the idea of bacterial proteases modulating host proteins and peptides is gaining traction.

Interestingly, the *Enterococcus* genus has been linked to obesity in children and adolescents, as well as to mice consuming a western diet (21, 22). While not characterized in this study, we also identified several *Clostridium* isolates whose supernatants are capable of decreasing GLP-1 levels in vitro. We suspect this activity is also the result of a secreted protease cleaving GLP-1, supporting the idea that the microbiome produces a suite of proteases capable of interfering with host metabolism. Although *E. faecalis* and *Clostridium* have been implicated in infection, but they can also behave as commensal organisms and often live inconspicuously in our GI tracts. It is important to understand all the interactions of these organisms with their host, not just the overt pathogenic functions.

In summary, the results of this study demonstrate that GelE, a recognized virulence factor of *E. faecalis*, can degrade the human GI hormone GLP-1, among other metabolic substrates. The degradation of GLP-1 likely occurs by slight damage to the intestinal epithelium, allowing GelE to translocate across the epithelial layer and access GLP-1. While it would be reckless to assume this activity is an etiology of metabolic disease, we do believe that interference with natural GLP-1 signaling by microbial degradation of GLP-1 could be a contributing factor to the development of disease. An important next step for this work is to assess the contribution of GelE to intestinal barrier permeability and the development of metabolic syndrome in vivo. Finally, this study adds a novel mechanism of action to the ever-growing list of host-microbe interactions.

## Materials and methods

### Bacterial strain isolation

Bacterial strains for screening were isolated previously from fecal, breast milk, and ileum biopsy samples (8). Genetic mutant strains of *Enterococcus faecalis* (OG1RF, TX5266 (*fsrB*), TX5264 (*gelE*), TX5243 (*sprE*), TX5128 (*gelE;sprE*)) were generously gifted from the Danielle A. Garsin Laboratory (University of Texas, Health Science Center) (10). *E. faecalis* S613 was generously gifted from the Cesar A. Arias Laboratory (University of Texas, Heath Science Center) (11).

### Bacterial growth and preparation of cell-free supernatants

Bacterial isolates were streaked from frozen glycerol stocks onto GM17 agar plates and incubated anaerobically overnight at 37°C. One colony was inoculated into 5 mL of GM17 broth and incubated overnight at 37°C followed by one more subculture into GM17 broth, and incubation overnight at 37°C. Once grown, bacterial cultures were centrifuged at 5000 x g for 20 min. Supernatants were collected and lyophilized (Labconco Freezone), followed by storage at −80°C until used for subsequent assays.

### 16S rRNA gene sequencing of isolates

To identify the bacterial isolates, bacteria were streaked on GM17 (M17 + 0.5% (w/v) glucose) agar plates from frozen glycerol stocks and incubated at 37°C for 24-48 h. Bacterial colony mass was then resuspended in 800 μL of sterile water and transferred to sterile bead beating tubes and homogenized for 2 min in a mini-beadbeater-96 (Biospec Products). Tubes were centrifuged at 8000 xg for 30 sec and supernatants were used for 16S rRNA gene PCR amplification with Phusion High-Fidelity DNA Polymerase (New England Biolabs) in a 20 μL reaction according to the manufacturer’s protocol, with sequencing primers 8F and 1492R. The amplification cycle consisted of an initial denaturation at 98°C for 30 sec, followed by 26 cycles of 10 sec at 98°C, 20 sec at 51°C, and 1 min at 72°C. Amplification was verified by agarose gel electrophoresis. For sample cleanup, DNA was treated with Exo-SAP-IT (ThermoFisher) and incubated at 37°C for 15 min followed by a 15 min incubation at 80°C to inactivate the enzyme. The product was cooled and sent to Genewiz for sequencing according to company protocol. Upon return of sequencing data, sequences were compared to the NCBI BLAST database.

### Screening for GLP-1 stimulatory activity using NCI H716 cells

NCI H716 (American Type Culture Collection (ATCC) CCL-251) cells were grown in Roswell Park Memorial Institute (RPMI, ATCC) medium supplemented with 10% (v/v) heat inactivated newborn calf serum (NBCS, Gibco). Cultures were maintained at a concentration of 2-8 × 10^5^ cells/mL and used at passages 15-40 for cell studies. For cell studies, 96-well plates were coated with 100 μL of 10 mg/mL Matrigel (BD Biosciences) for 2 h at room temperature. Following coating, NCI H716 cells were seeded at a concentration of 1 × 10^5^ cells/well in Dulbecco’s Modified Eagle’s Medium (DMEM) supplemented with 10% (v/v) NBCS, as determined by trypan blue staining using a hemocytometer. Two days later, lyophilized bacterial supernatants were resuspended in Krebs buffer (Sigma) containing bovine serum albumin (BSA, 0.2% w/v) and bovine bile (0.03% w/v) and incubated on the NCI H716 cells at 37°C with 5% CO2. 4-phorbol 12 myristate 13-acetate (PMA, 2 μM) was used as a positive control as it is a potent stimulator of GLP-1 secretion through activation of protein kinase C (PKC). Following a 2h incubation, supernatants were collected and analyzed for total GLP-1 levels by ELISA (Millipore Sigma) according to the manufacturer’s protocol. Cell viability was monitored using PrestoBlue Cell Viability Reagent (ThermoFisher Scientific) following the manufacturer’s instructions.

### Characterization studies (TPEN, heat, size)

For metalloprotease inhibitor studies, *E. faecalis* strains were subcultured 0.1% (v/v) from an overnight culture into GM17 containing the indicated concentration of N,N,N’,N’-tetrakis(2-pyridylmethyl)ethane-1,2-diamine (TPEN). Cultures were grown overnight, supernatants were collected as described above, and incubated with 500 pM GLP-1 (GenScript) for 4 h at room temperature, followed by storage at −80°C until ready for GLP-1 quantification by ELISA.

For heat treatment studies, bacterial supernatants were heated to 90°C for 30 min. Samples were then used in an NCI H716 cell assay as described above, followed by storage at −80°C until ready for GLP-1 quantification by ELISA.

For size fractionation studies, bacterial supernatants were separated by size using centrifugal filter units (Amicon) and centrifuged as described above. Samples were then used in an NCI H716 cell assay as described above, followed by storage at −80°C until ready for GLP-1 quantification by ELISA.

### Protease knock-out studies

Supernatants from *E. faecalis* protease mutants and controls (OG1RF, TX5266 (*fsrB*), TX5264 (*gelE*), TX5243 (*sprE*), TX5128 (*gelE;sprE*)) were collected from an overnight culture grown in GM17 aerobically at 37°C by centrifugation as described above. Supernatants were incubated with 500 pM GLP-1 (Tocris or GenScript) for 4 h at room temperature, followed by storage at −80°C until ready for GLP-1 quantification by ELISA, as described above.

### RNA collection and quantitation of gelE expression

*E. faecalis* were subcultured 1% (v/v) from an overnight culture into GM17. After 5 h incubation, cells were collected by centrifugation, resuspended in RNALater solution (Invitrogen), and stored at −80°C. Cells were washed in 1X PBS, resuspended in 1 mL RTL buffer (Qiagen RNeasy Kit) and lysed by bead beating (2 × 1 min) at 4°C followed by RNA extraction according to the manufacturer’s instructions. cDNA was synthesized using Superscript III reverse transcriptase (Invitrogen) following the manufacturer’s recommended protocol. Quantitative PCR reactions were performed using Power SYBR Green Master Mix (Applied Biosystems) with either *E. faecalis* 16s RNA (f: 5’-CCGAGTGCTTGCACTCAATTGG-3’, r: 5’-CTCTTATGCCATGCGGCATAAAC-3’) or gelE (f: 5’-CGGAACATACTGCCGGTTTAGA-3’, r: 5’-TGGATTAGATGCACCCGAAAT-3’) specific primers (Wang et al. 2011). The expression of gelE was normalized to that of the 16S RNA and the data were analyzed using the 2 ^−Δ ΔCT^.

### Cell culture growth and assays of T84 monolayers

Growth and assays of T84 cells were performed by methods described previously, with slight modifications (Zeng 2004, Hopper 2000). T84 human colonic epithelial cells (ATCC CCL-248) were propagated in tissue culture-treated T75 flasks (CELLSTAR) as indicated by ATCC. When between 90-100% confluency, T84 cells were treated with 0.25% trypsin and plated onto 24-well, 3.0 μm polycarbonate membrane transwell filters (Costar, 3415) at a density of 8 × 10^4^ cells/well. The electrical resistance of the monolayer was monitored over the course of 2-3 weeks, and monolayers with a transepithelial electrical resistance (TEER, Millipore Millicell ERS-2) >800 Ω/cm^2^ were used for GLP-1 cleavage assays.

To prepare bacteria, on the day of the assay *E. faecalis* strains were subcultured 1% (v/v) from an overnight culture into GM17 and grown for 5 h as described above. Whole culture (cells + supernatant) samples were diluted in DMEM/F12 (Gibco) to a concentration of 1 × 10^7^ CFU/mL, and supernatant samples were diluted 50% in DMEM/F12. All samples were neutralized to a pH of 6.8-7 using 3M NaOH.

Once prepared, the TEER was measured for each monolayer, followed by 100 μL of *E. faecalis* sample added to the upper chamber of the transwell, and 500 μL of tissue culture media containing 500 pM GLP-1 (GenScript) added to the lower chamber. After a 16 h incubation at 37°C in 5% CO2, 200 μL of media from the lower chamber was removed and stored at −80°C until ready for GLP-1 quantification by ELISA, as described above. A final TEER measurement was taken for each monolayer.

### Luminex

A Milliplex Multiplex assay was performed using a Metabolic Luminex kit (Millipore Sigma) to measure total GLP-1, glucagon, Gastric Inhibitory Peptide/Glucose Insuliotropic Peptide (GIP), leptin, Peptide YY (PYY), pancreatic peptide, insulin, monocyte chemoattractant protein-1 (MCP-1), interleukin-6 (IL-6), and tumor necrosis factor alpha (TNFa). Bacterial supernatants were collected from an overnight culture by centrifugation as described above, diluted in bacterial culture media as indicated, and incubated with recombinant protein (GenScript, Sigma, Tocris) of each analyte for 4 h at room temperature, followed by storage at −80°C until ready for analyte quantification. The Milliplex Multiplex assay was run according to the protocol provided by the manufacturer.

### Statistical analysis

Statistical analyses were performed using GraphPad Prism version 8.0 (San Diego, CA, USA). Experimental results are expressed as means ± standard deviation. Statistical significance was set at p < 0.05. One-way statistical comparisons were carried out using one-way analysis of variance (ANOVA), followed by multiple comparisons of the means using Tukey’s post-hoc analysis, for the GLP-1 secretion experiments in NCI-H716 cells, protease knock-out mutation assay, and degradation preference of GelE using a Luminex assay (undiluted supernatants). Two-way ANOVA analysis, followed by multiple comparisons of the means using Tukey’s post-hoc analysis, was performed for the degradation preference of GelE using a Luminex assay (diluted supernatants) and T84 cell experiments.

## Acknowledgements

We would like to acknowledge the Functional Genomics and Microbiome Core of the Texas Medical Center Digestive Diseases Center at Baylor College of Medicine for the help and guidance performing the Luminex assays. Also, the laboratories of Danielle A. Garsin and Cesar A. Arias at the University of Texas, Heath Science Center for the generous donations of *E. faecalis* strains. This work was supported in part by Baylor College of Medicine seed funding. CTD acknowledges funding in the form of a postdoctoral fellowship from the Fonds de recherche santé Québec.

## Supplemental figures

**S1.**
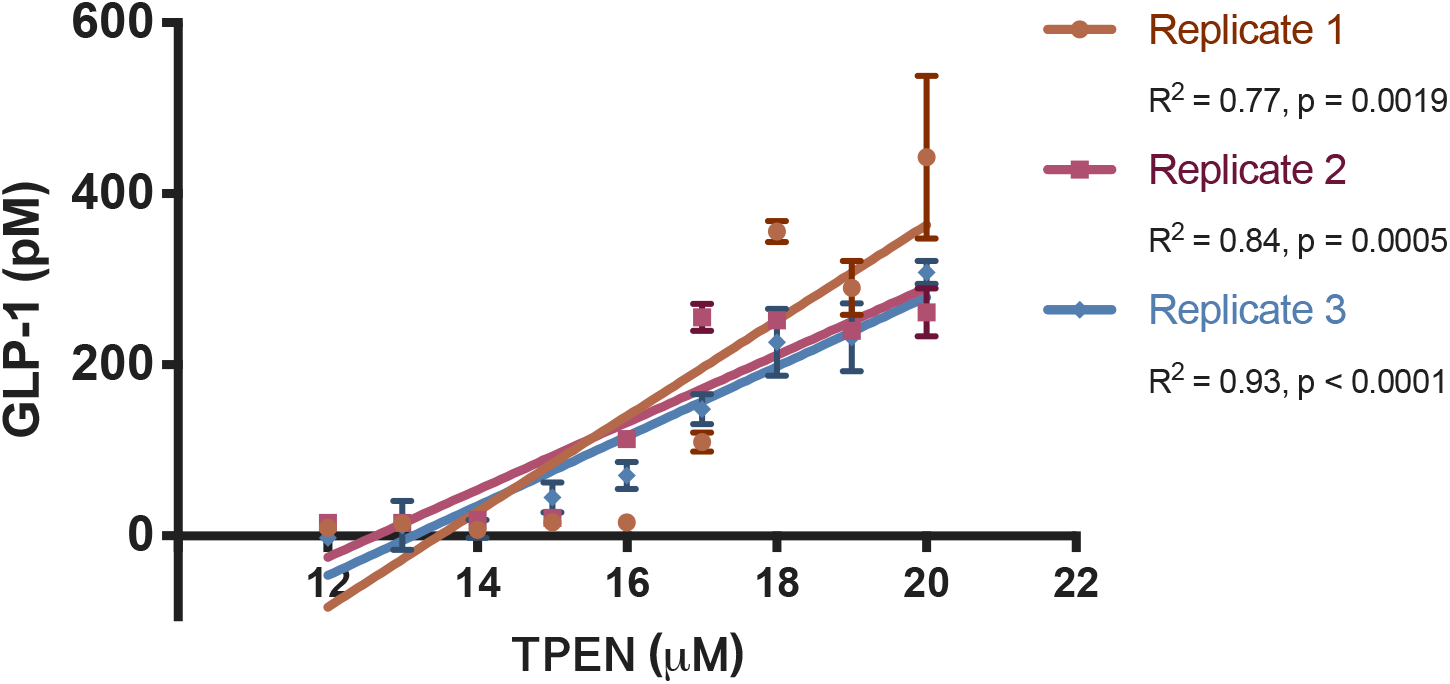
GLP-1 inhibitory activity is sensitive to TPEN. Supernatnats from *E. faecais* 30054A grown overnight with increasing amounts of metalloprotease inhibitor TPEN (higher specificity for zinc) result in a reduction of GLP-1 degradation when incubated with recombinant GLP-1, in a dose-dependent manner.

**S2.**
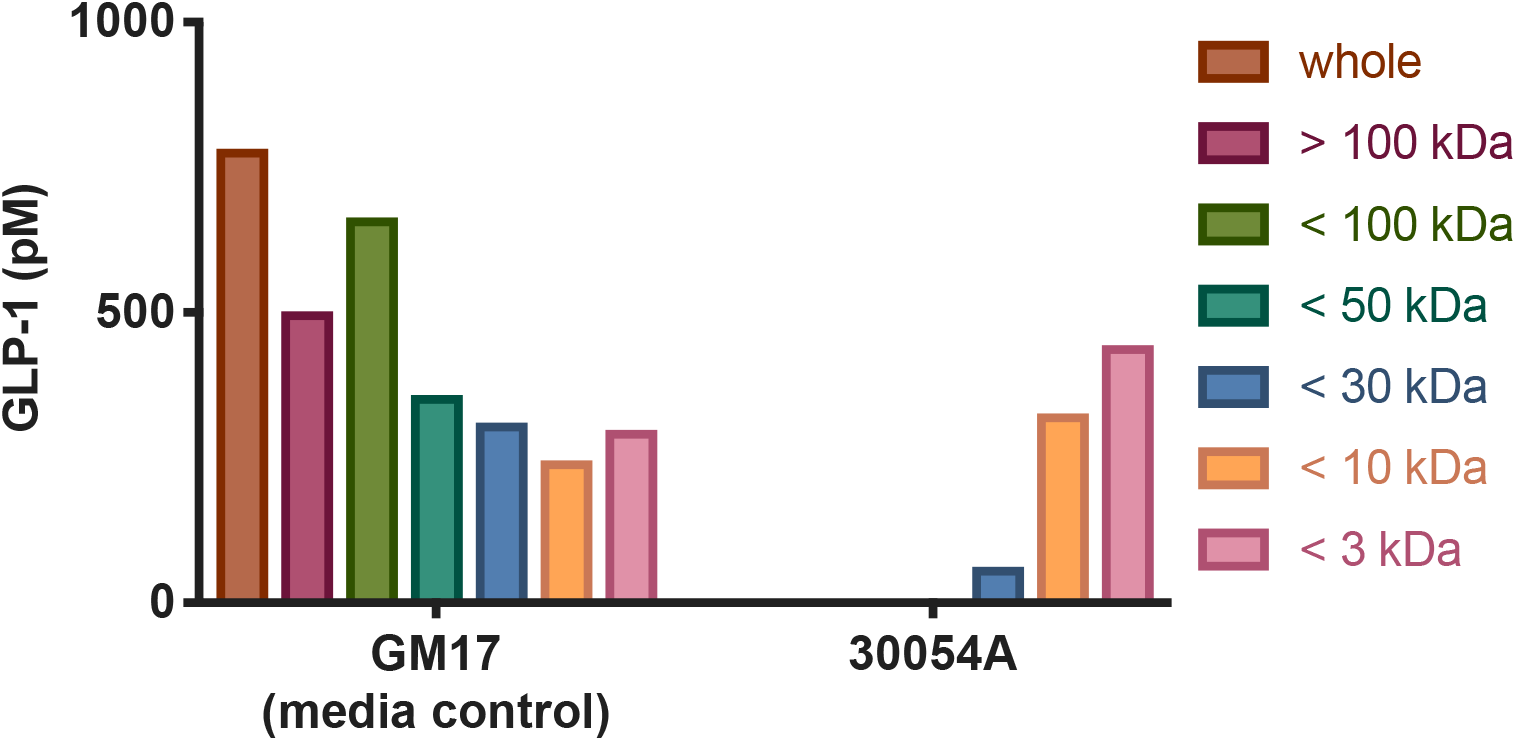
GLP-1 inhibitory activity is contained in the >30 kDa size fraction. Supernatants from *E. faecalis* 30054A were fractioned using centrifugal filters of various kDa cutoffs, and then incubated on NCI H716 cells. GLP-1 inhibitory activity is maintained in all fractions >30 kDa.

**S3.**
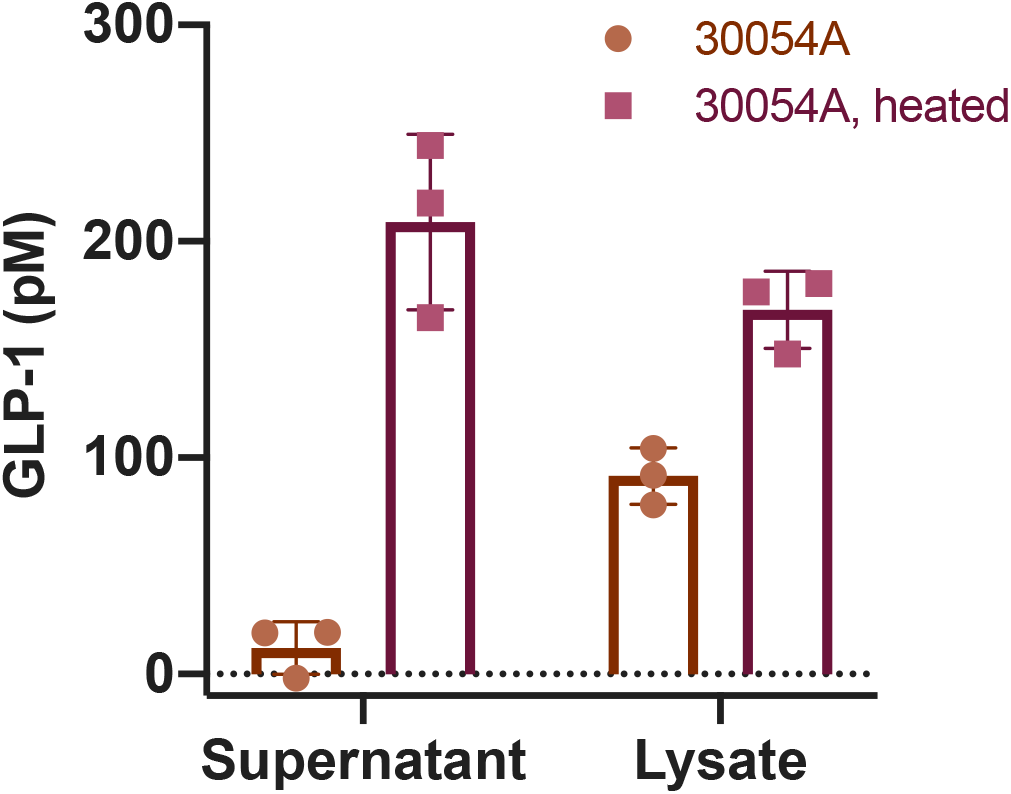
GLP-1 inhibitory activity is lost with heat treatment. Supernatants and whole cell lysates from *E. faecalis* 30054A treated to 30 min at 100°C result in a reduction of GLP-1 inhibitory activity when incubated with NCI H716 cells. GLP-1 inhibitory activity is primarily contained in the bacterial supernatant as opposed to the cellular lysate.

## Bibliography

1. Gilbert JA, Blaser MJ, Caporaso JG, Jansson JK, Lynch SV, Knight R. 2018. Current understanding of the human microbiome. Nat Med 24:392–400.

2. Lynch SV, Pedersen O. 2016. The Human Intestinal Microbiome in Health and Disease. New England Journal of Medicine 375:2369–2379.

3. Bäckhed F, Ding H, Wang T, Hooper LV, Koh GY, Nagy A, Semenkovich CF, Gordon JI. 2004. The gut microbiota as an environmental factor that regulates fat storage. Proceedings of the National Academy of Sciences of the United States of America 101:15718.

4. Cani PD, Neyrinck AM, Maton N, Delzenne NM. 2005. Oligofructose Promotes Satiety in Rats Fed a High-Fat Diet: Involvement of Glucagon-Like Peptide-1. Obesity Research 13:1000–1007.

5. Everard A, Belzer C, Geurts L, Ouwerkerk JP, Druart C, Bindels LB, Guiot Y, Derrien M, Muccioli GG, Delzenne NM, de Vos WM, Cani PD. 2013. Cross-talk between *Akkermansia muciniphila* and intestinal epithelium controls diet-induced obesity. Proceedings of the National Academy of Sciences 110:9066.

6. Plovier H, Everard A, Druart C, Depommier C, Van Hul M, Geurts L, Chilloux J, Ottman N, Duparc T, Lichtenstein L, Myridakis A, Delzenne NM, Klievink J, Bhattacharjee A, van der Ark KC, Aalvink S, Martinez LO, Dumas ME, Maiter D, Loumaye A, Hermans MP, Thissen JP, Belzer C, de Vos WM, Cani PD. 2017. A purified membrane protein from Akkermansia muciniphila or the pasteurized bacterium improves metabolism in obese and diabetic mice. Nat Med 23:107–113.

7. Reimer RA, Darimont C, Gremlich S, Nicolas-Metral V, Ruegg UT, Mace K. 2001. A human cellular model for studying the regulation of glucagon-like peptide-1 secretion. Endocrinology 142:4522–4528.

8. Tomaro-Duchesneau C, LeValley SL, Röth D, Sun L, Horrigan FT, Kalkum M, Hyser JM, Britton RA. 2018. Discovery of a bacterial peptide as a modulator of GLP-1 and metabolic disease. bioRxiv doi:10.1101/379503:379503.

9. Podbielski A, Kreikemeyer B. 2004. Cell density--dependent regulation: basic principles and effects on the virulence of Gram-positive cocci. Int J Infect Dis 8:81–95.

10. Cruz MR, Graham CE, Gagliano BC, Lorenz MC, Garsin DA. 2013. Enterococcus faecalis inhibits hyphal morphogenesis and virulence of Candida albicans. Infect Immun 81:189–200.

11. Arias CA, Panesso D, McGrath DM, Qin X, Mojica MF, Miller C, Diaz L, Tran TT, Rincon S, Barbu EM, Reyes J, Roh JH, Lobos E, Sodergren E, Pasqualini R, Arap W, Quinn JP, Shamoo Y, Murray BE, Weinstock GM. 2011. Genetic Basis for In Vivo Daptomycin Resistance in Enterococci. New England Journal of Medicine 365:892–900.

12. Makinen PL, Clewell DB, An F, Makinen KK. 1989. Purification and substrate specificity of a strongly hydrophobic extracellular metalloendopeptidase (“gelatinase”) from Streptococcus faecalis (strain 0G1-10). J Biol Chem 264:3325–3334.

13. Steck N, Hoffmann M, Sava IG, Kim SC, Hahne H, Tonkonogy SL, Mair K, Krueger D, Pruteanu M, Shanahan F, Vogelmann R, Schemann M, Kuster B, Sartor RB, Haller D. 2011. Enterococcus faecalis metalloprotease compromises epithelial barrier and contributes to intestinal inflammation. Gastroenterology 141:959–971.

14. Zeng J, Teng F, Murray BE. 2005. Gelatinase is important for translocation of Enterococcus faecalis across polarized human enterocyte-like T84 cells. Infect Immun 73:1606–1612.

15. Maharshak N, Huh EY, Paiboonrungruang C, Shanahan M, Thurlow L, Herzog J, Djukic Z, Orlando R, Pawlinski R, Ellermann M, Borst L, Patel S, Dotan I, Sartor RB, Carroll IM. 2015. Enterococcus faecalis Gelatinase Mediates Intestinal Permeability via Protease-Activated Receptor 2. Infect Immun 83:2762–2770.

16. Waters CM, Antiporta MH, Murray BE, Dunny GM. 2003. Role of the Enterococcus faecalis GelE protease in determination of cellular chain length, supernatant pheromone levels, and degradation of fibrin and misfolded surface proteins. J Bacteriol 185:3613–3623.

17. Hancock LE, Perego M. 2004. The *Enterococcus faecalis fsr* Two-Component System Controls Biofilm Development through Production of Gelatinase. Journal of Bacteriology 186:5629.

18. Mohamed JA, Huang W, Nallapareddy SR, Teng F, Murray BE. 2004. Influence of Origin of Isolates, Especially Endocarditis Isolates, and Various Genes on Biofilm Formation by *Enterococcus faecalis*. Infection and Immunity 72:3658.

19. Shaked H, Carmeli Y, Schwartz D, Siegman-Igra Y. 2006. Enterococcal bacteraemia: epidemiological, microbiological, clinical and prognostic characteristics, and the impact of high level gentamicin resistance. Scand J Infect Dis 38:995–1000.

20. Park SY, Shin YP, Kim CH, Park HJ, Seong YS, Kim BS, Seo SJ, Lee IH. 2008. Immune Evasion of *Enterococcus faecalis* by an Extracellular Gelatinase That Cleaves C3 and iC3b. The Journal of Immunology 181:6328.

21. Hou Y-P, He Q-Q, Ouyang H-M, Peng H-S, Wang Q, Li J, Lv X-F, Zheng Y-N, Li S-C, Liu H-L, Yin A-H. 2017. Human Gut Microbiota Associated with Obesity in Chinese Children and Adolescents. BioMed research international 2017:7585989–7585989.

22. Turnbaugh PJ, Ridaura VK, Faith JJ, Rey FE, Knight R, Gordon JI. 2009. The effect of diet on the human gut microbiome: a metagenomic analysis in humanized gnotobiotic mice. Science translational medicine 1:6ra14–16ra14.

23. Shaked H, Carmeli Y, Schwartz D, Siegman-Igra Y. 2006. Enterococcal bacteraemia: Epidemiological, microbiological, clinical and prognostic characteristics, and the impact of high level gentamicin resistance. Scandinavian Journal of Infectious Diseases 38:995–1000.

24. Zietek T, Rath E. 2016. Inflammation Meets Metabolic Disease: Gut Feeling Mediated by GLP-1. Frontiers in Immunology 7.

25. Olivares M, Schüppel V, Hassan AM, Beaumont M, Neyrinck AM, Bindels LB, Benítez-Páez A, Sanz Y, Haller D, Holzer P, Delzenne NM. 2018. The Potential Role of the Dipeptidyl Peptidase-4-Like Activity From the Gut Microbiota on the Host Health. Frontiers in Microbiology 9:1900.

